# A Clinical Phenotyping Algorithm to Identify Cases of Chronic Obstructive Pulmonary Disease in Electronic Health Records

**DOI:** 10.1101/716779

**Authors:** Victoria L. Martucci, Nancy Liu, V. Eric Kerchberger, Travis J. Osterman, Eric Torstenson, Bradley Richmond, Melinda C. Aldrich

## Abstract

**Rationale:** Chronic obstructive pulmonary disease (COPD) is a leading cause of mortality in the United States. Electronic health records provide large-scale healthcare data for clinical research, but have been underutilized in COPD research due to challenges identifying these individuals, especially in the absence of pulmonary function testing data.

**Objectives:** To develop an algorithm to electronically phenotype individuals with COPD at a large tertiary care center.

**Methods:** We identified individuals over 45 years of age at last clinic visit within Vanderbilt University Medical Center electronic health records. We tested phenotyping algorithms using combinations of both structured and unstructured text and examined the clinical characteristics of the resulting case sets.

**Measurement and Main Results:** A simple algorithm consisting of 3 International Classification of Disease codes for COPD achieved a sensitivity of 97.6%, a specificity of 76.0%, a positive predictive value of 57.1%, and a negative predictive value of 99.0%. A more complex algorithm consisting of both billing codes and a mention of oxygen on the problem list that achieved a positive predictive value of 86.5%. However, the association of known risk factors with chronic obstructive pulmonary disease was consistent in both algorithm sets, suggesting a simple code-only algorithm may suffice for many research applications.

**Conclusions:** Simple code-only phenotyping algorithms for chronic obstructive pulmonary disease can identify case populations with epidemiologic and genetic profiles consistent with published literature. Implementation of this phenotyping algorithm will expand opportunities for clinical research and pragmatic trials for COPD.

## Introduction

Chronic obstructive pulmonary disease (COPD) is a leading cause of death globally.(1) Within the United States, approximately 6.3% of adults live with COPD, contributing to $32.1 billion in health care costs annually.(2, 3) Individuals with COPD have high rates of hospitalization, health care utilization, and all-cause mortality, as well as increased mortality from comorbid conditions such as cardiovascular disease, kidney disease, and lung cancer.(4–7) COPD is characterized by small airway narrowing and obliteration and emphysematous lung destruction, which together lead to progressive obstruction in expiratory airflow.(8) The clinical gold standard for a COPD diagnosis is demonstration of irreversible airflow limitation assessed via pulmonary function testing (PFT).(8, 9) However, routine screening for COPD is not recommended, particularly among asymptomatic patients,(10) and PFTs are underutilized in clinical settings.(11, 12) Discovering a reliable means to identify COPD cases in electronic health records (EHRs) in the absence of PFTs could greatly facilitate advances in COPD clinical research. To date, most research studies have relied on well-curated cohorts to further our understanding of COPD risks and outcomes.(13–17) These observational cohorts are costly and time-consuming to develop, hampering rapid advances in research. EHR provide a valuable tool to expand and potentially accelerate clinical research of COPD.(18) EHR also allow unique opportunities for pragmatic clinical trials, with an unprecedented depth of information that can be used to rapidly and cost-effectively identify trial participants, study medical interventions and outcomes in real-world settings, and prioritize interventions for testing in randomized controlled trials.(19–22) The development of a robust and portable algorithm to identify COPD cases in the absence of PFTs is needed to enable genetics research leveraging biobanks and pragmatic clinical trials for COPD.

Relatively few attempts have been made to develop such an algorithm for COPD. Himes et al. developed an algorithm to predict COPD in asthma patients, but the generalizability of this algorithm to COPD not associated with asthma is unclear.(23) Several algorithms to identify COPD within primary care and administrative claims databases have been developed.(24–31) The performance of these algorithms varies greatly, with positive predictive values (PPV) ranging from 36.7% to 94%.(24, 26, 27, 29–31) However, algorithms with higher PPVs had lower sensitivities, meaning many cases of COPD were missed. These algorithms were predominantly developed in primary care populations in Canada and Europe. Therefore, the generalizability to hospital-based populations in the United States is unclear. To our knowledge, the only phenotyping algorithm for COPD in United States hospital-based clinical populations was used in a recent genetic association study by Wain et al.,(32) yet the algorithm performance was not reported.

Vanderbilt University Medical Center (VUMC) has a well-characterized EHR of clinical data captured through routine care. We used de-identified data from the VUMC clinical population to develop and evaluate EHR-based COPD phenotyping algorithms. We also characterized the COPD cases with regards to established risk factors, candidate genetic variants, and comorbidities.

## Methods

### Synthetic Derivative

We used clinical data from the Vanderbilt Synthetic Derivative (SD), a de-identified version of the VUMC EHR, containing data on over 2.1 million adult patients and over 1 billion unique observations dating back to the 1980s.(33) Details regarding SD development have been previously published.(33) Extractable PFT data have been available since 2011. The Vanderbilt University Institutional Review Board approved this study.

Demographic data, International Classification of Disease (ICD) 9 and ICD-10 codes, and PFTs were obtained from structured fields in the SD. Quality control was implemented to remove individuals with record lengths (defined as the number of days from the first clinical encounter and the most recent clinical encounter) longer than their age. Natural language processing mined additional text for medications for treatment of COPD; radiology report mentions of emphysema (excluding subcutaneous emphysema); and mentions of COPD, emphysema, chronic bronchitis, cough, shortness of breath, or oxygen use on the problem list (Table S1). We implemented negation for radiology reports using PyConTextNLP, a Python algorithm that considers the context around keywords, and a previously curated list of negation terms.(34, 35) Smoking information was collected from unstructured clinical notes. Due to current lack of granularity in patient smoking behaviors in the clinical record,(36) we simplified smoking information to a dichotomous variable of ever versus never smoker.

### Algorithm Development and Validation

The study population consisted of adults over 45 years of age at last clinic visit who visited VUMC prior to March 8, 2019. We identified individuals with available PFT data and randomly sampled 200 patients for our development set (Figure 1). PFTs were used as the gold standard, with cases defined as individuals with a forced expiratory volume in one second (FEV_1_) to forced vital capacity (FVC) < 0.7 after bronchodilator administration. We developed a series of algorithms combining COPD ICD codes and additional clinical data (Table S1). To internally validate our algorithm performance, we tested them in an independent random sample of 200 records. Stratified random sampling was used to select 100 individuals with two or more COPD ICD-9 or ICD-10 codes and 100 individuals with fewer than two COPD ICD codes. Gold standard chart review was performed by two independent reviewers (VLM and VEK) with clinical training. Discrepancies were adjudicated by a pulmonary physician (BR). Kappa statistics were calculated to determine agreement between reviewers.(37) A second internal validation was performed using all individuals with available PFTs, excluding the 200 records used in the development set (N = 13,858).

**Figure 1.**
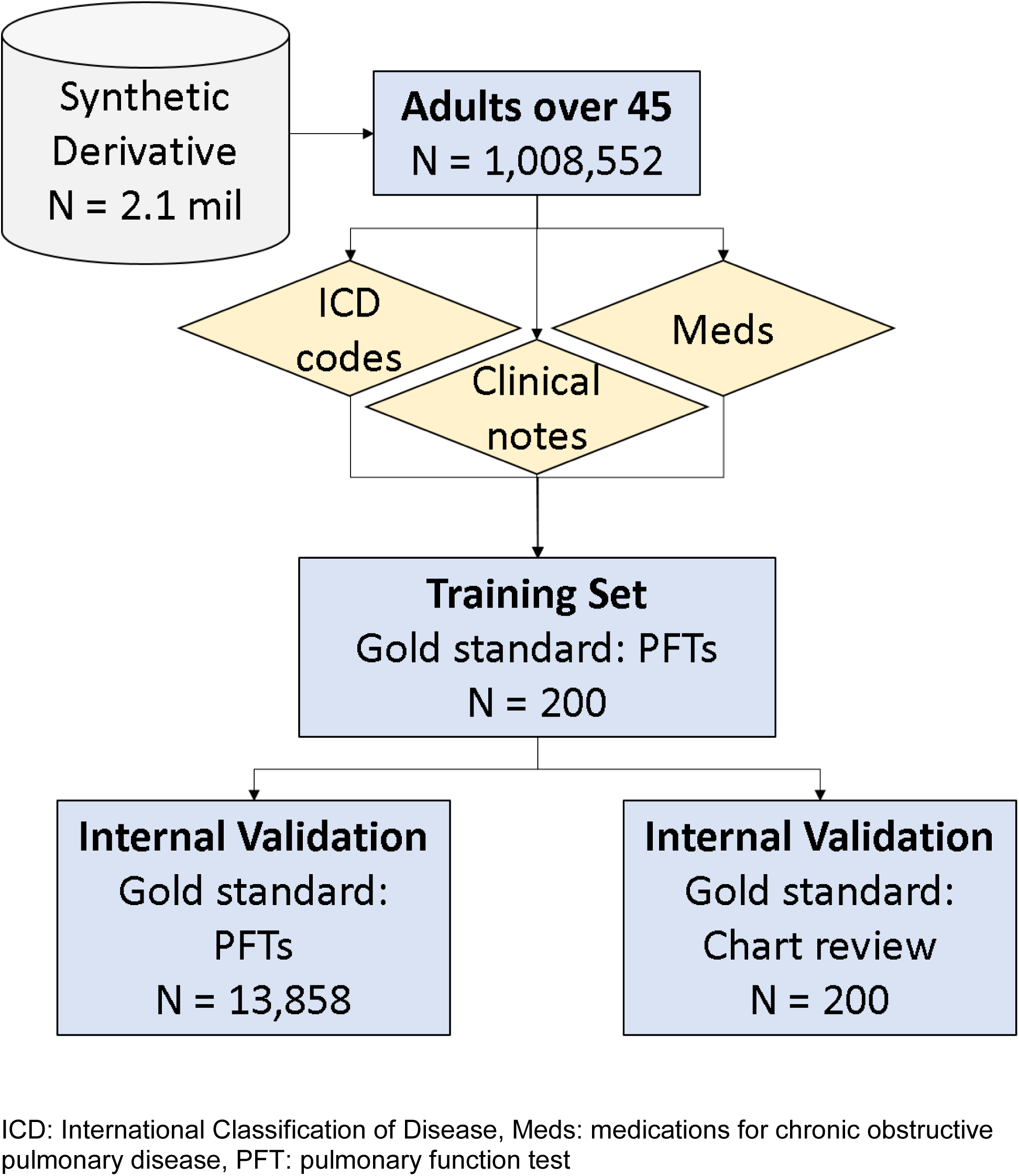
Study design for algorithm development and Vanderbilt Synthetic Derivative sample size as of March 2019.

### Applications of Phenotyping Algorithm

We applied the algorithms to the entire SD population over 45 years of age at last visit to calculate the number of cases, controls, indeterminates, and excluded individuals. To ensure cases and controls had similar opportunity for eligibility, we selected a study population with a minimum record length of 180 days (6 months). We then examined the relationship between known risk factors (age, sex, and smoking history) and algorithm-defined COPD case-control status using logistic regression models.

To demonstrate potential applications of our algorithms, we explored comorbidity data within our algorithm-defined cases using phecodes. Phecodes are aggregated ICD-9 codes that condense similar disease entities into one code, reducing the number of phenotypes from over 14,000 separate ICD-9 codes to 1,645 phecode groups, and have been utilized in multiple EHR and genetic studies.(38–44) We also conducted genetic analyses using COPD-associated single nucleotide polymorphisms (SNPs) previously identified by Wain et al.(32) Genotyping data was obtained from BioVU, a Vanderbilt biobank with de-identified genotyping and clinical data.(33) Data from the Illumina MEGA-Ex array were subjected to quality control to remove individuals and SNPs with <98% call rates, SNPs with minor allele frequency <1%, and SNPs not in Hardy-Weinberg equilibrium (p-value threshold of 1×10^−6^). After quality control, genotyping data were available on 41,660 physician-reported white adults over 45 years of age. Principal component (PC) analysis was performed using EIGENSTRAT on a set of SNPs pruned for linkage disequilibrium.(45) A log-additive model was assumed for individual SNPs and logistic regression analyses were performed to examine the association between SNPs and COPD, adjusting for age at last clinic visit, age^2^, sex, height, and the first 10 PCs for ancestry using Plink v1.9.(46)

## Results

### Study Population

As of March 8, 2019, we identified 1,008,661 individuals age 45 or older at last clinic visit. Quality control removed 109 individuals, leaving 1,008,552 individuals. For our initial algorithm development, we randomly sampled 200 charts with PFT data. The median age at last clinic visit was 65 years, with a median record length of 8.6 years. The development set had roughly equal percentages of males (49.0%) and females (51.0%) and was largely of observer-reported European descent (82.0%). The majority were smokers (56.5%) (Table 1). Our first internal validation set consisted of 200 records that underwent chart review. The population characteristics of the chart review validation set were similar to those of the development set, although there were fewer smokers in this set (40.5%) (Table 1). The kappa statistic between clinical reviewers was 0.75, and 18 discrepancies were adjudicated by a third reviewer. Our second internal validation set consisted of all individuals with available PFTs, excluding records used for development. The PFT validation set had similar characteristics to the chart review validation set, although individuals in this set had a longer record length (median 8.8 years) and higher smoking percentage (55.0%) than the chart review set or the overall SD (Table 1).

**Table 1.** Demographic characteristics of development set, internal validation sets, and all adults over age 45 years in the Synthetic Derivative, as of March 2019.

| **Variable** | **Development Set**<br>**N = 200** | **Internal Validation**<br>**Set (Chart Review)**<br>**N = 200** | **Internal Validation**<br>**Set (PFTs)**<br>**N = 13,858** | **SD over age 45**<br>**N = 1,008,552** |
| --- | --- | --- | --- | --- |
| Median age at last clinic visit, years (IQR) | 65<br>(58-73) | 64<br>(58-74) | 65<br>(57-73) | 61<br>(53-71) |
| Female, N (%) | 102 (51.0) | 107 (53.5) | 7,306 (52.7) | 545,271 (54.1) |
| Race, N (%) |  |  |  |  |
| White | 164 (82.0) | 156 (78.0) | 12,010 (86.7) | 702,679 (69.7) |
| Black | 28 (14.0) | 17 (8.5) | 1,374 (9.9) | 75,554 (7.5) |
| Other | 7 (3.5) | 2 (1.0%) | 180 (1.3) | 9,007 (0.9) |
| Median record length, years (IQR) | 8.6<br>(4.0-15.4) | 4.3<br>(0.9-10.5) | 8.8<br>(3.7-15.0) | 1.9<br>(0.1-8.2) |
| Ever smoker, N (%) | 113 (56.5) | 81 (40.5) | 7,619 (55.0) | 190,986 (18.9) |
| COPD prevalence, N (%) |  |  |  |  |
| Algorithm-based | 66 (33.0) | 70 (35.0) | 3,834 (27.7) | 34,513 (3.4) |
| PFT-based | 37 (18.5) | NA | 1,976 (14.3) | NA |

SD = synthetic derivative

### Algorithm development and performance in validation sets

We tested algorithms using different combinations of clinical text and COPD ICD codes in our development set (Table S1). Based on algorithm performance and complexity, we proceeded with two case algorithms and one control algorithm in the validation phase. The case and control definitions were as follows:

1. 3+ codes: A rule-based classifier that required three or more ICD codes for COPD (Figure 2)
2. Code + regex: A rule-based classifier that required at least 10 or more ICD codes for COPD *OR* presence of three to nine ICD codes for COPD *AND* a text mention of oxygen use on the problem list (Figure S1).
3. Controls: A rule-based classifier requiring no ICD codes for COPD *AND* no ICD codes for asthma *OR* idiopathic pulmonary fibrosis *OR* sarcoidosis (Figure 2).

**Figure 2.**
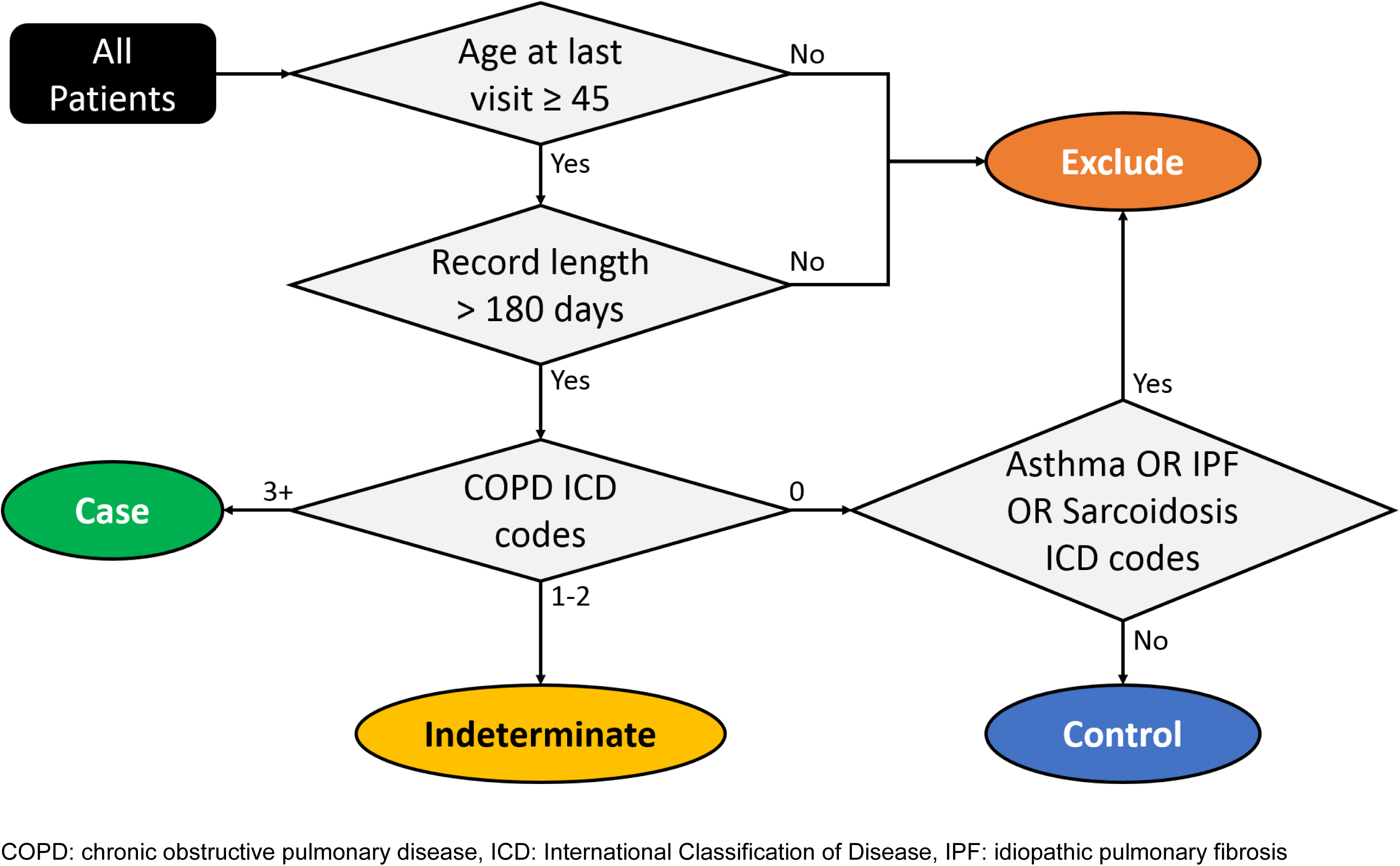
Phenotyping algorithm for chronic obstructive pulmonary disease.

The 3+ code algorithm had a sensitivity of 97.6%, specificity of 76.0%, PPV of 57.1%, NPV of 99.0%, and F-measure of 0.72 in the chart review validation set (Table 2). All calculated performance metrics were lower in the PFT validation set (Table 2). The code + regex algorithm had the highest specificity (95.0%), PPV (86.5%), and F-measure (0.91) in the chart review validation set (Table 2). This algorithm also had the best performance in terms of specificity (79.1%), PPV (39.8%), and F-measure (0.53) in the PFT validation set (Table 2).

**Table 2.**
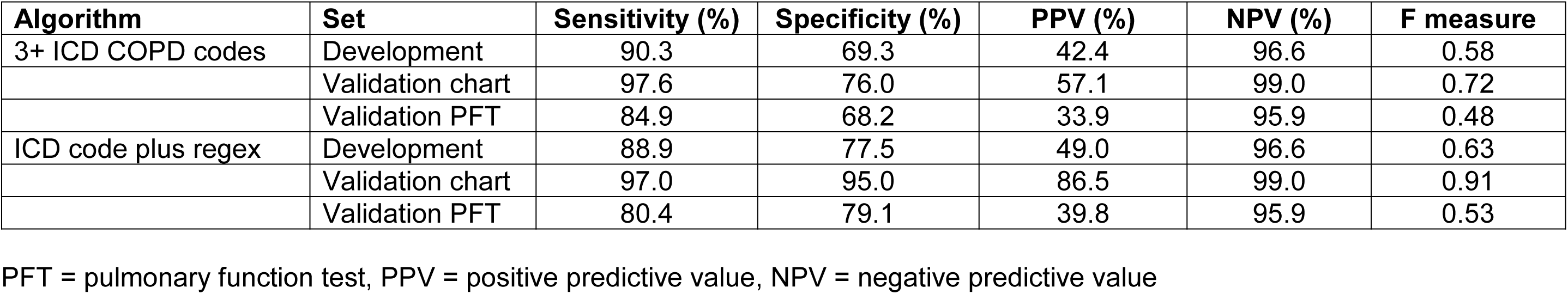
Clinical validity measures for phenotyping algorithms in development, validation, and PFT sets.

### Application of algorithms to EHR dataset

To demonstrate the utility of our algorithms within a large EHR database, we applied them to the entire adult SD population over 45 years of age at last visit with record lengths longer than 6 months (180 days, N = 623,986). The 3+ code-only algorithm identified 28,520 COPD cases (5.0% of adults meeting inclusion criteria) and 544,056 controls (95.0%) (Table S2). There were 23,091 individuals labeled as indeterminates due to too few COPD ICD codes and 28,319 individuals excluded from the control set due to the presence of asthma, sarcoidosis, or idiopathic pulmonary fibrosis codes. Using the code + regex algorithm decreased the case number to 12,622 (2.3%) (Table S2). Exclusively using PFT-defined COPD identified 2,015 individuals (14.3%) who met the GOLD definition for COPD (post-bronchodilator FEV_1_/FVC < 0.7) (Table S2).(8)

We compared the case sets identified by the 3+ code algorithm, the code + regex algorithm, and the gold standard PFT definition. The age, sex, and race distributions were similar in all three groups (Table S3). PFT-defined cases had a higher prevalence of ever smokers (1,464, 78.2%) compared to 3+ ICD code COPD cases (15,645, 54.9%) and code + regex COPD cases (7,548, 59.8%). The median record length was longer in PFT-defined cases (10.3 years) than in 3+ ICD code cases (7.3 years) and code + regex cases (8.3 years). We also looked at the overlap in sample size between cases identified by the 3+ code algorithm and those identified by PFTs. There were 1,265 cases present in both sets (Figure 3). The remaining 607 PFT cases (32.4% of all PFT cases with record lengths longer than 6 months) were not identified by our algorithm, and 27,255 algorithm cases were not identified by PFTs. We compared FEV_1_ percent predicted and FEV_1_/FVC measurements in PFT cases with 3+ ICD codes and PFT cases with fewer than three ICD codes. In both pre- and post-bronchodilator measures, the median FEV_1_ percent predicted and FEV_1_/FVC were lower among PFT cases with 3+ COPD ICD codes than those with fewer codes (Figure S2).

**Figure 3.**
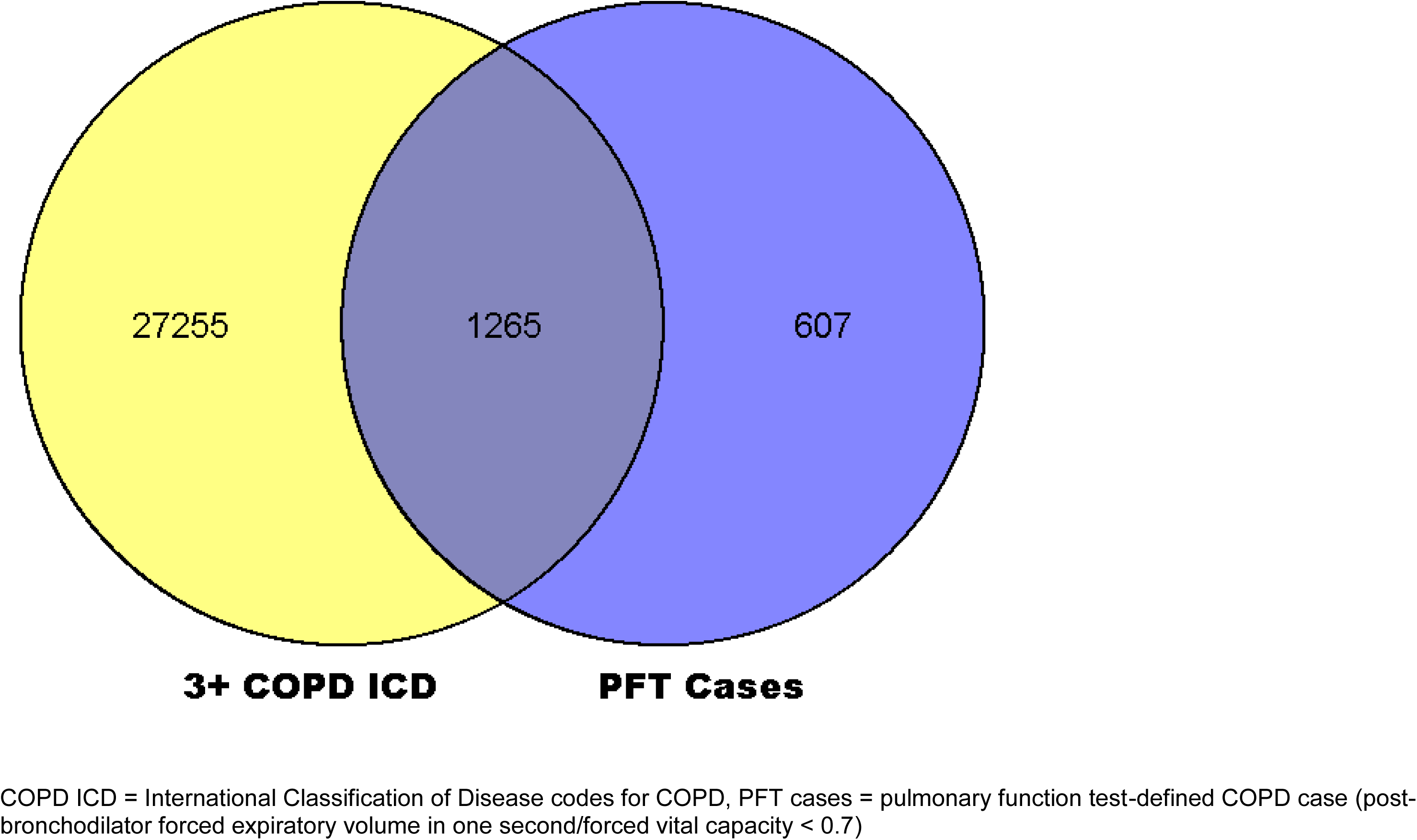
Overlap in COPD case definitions among COPD cases over age 45 years in the Synthetic Derivative.

### Confirmation of known clinical risk factors in algorithm COPD cases

To further inform our algorithm choice, we compared the odds ratios (OR) examining the associations between COPD status and known clinical risk factors in each algorithm-defined population. Using the 3+ code algorithm, the OR for COPD associated with age at last clinic visit was 1.04, with male sex was 1.40, and with ever smoking was 9.22. The code + regex algorithm and PFT-based definition all showed similar OR for age, sex, and smoking (Table S4). Since the associations did not differ meaningfully between the different algorithms tested, we chose to focus on the 3+ code algorithm due to its simplicity and larger case sample size.

### Clinical and genetic analyses using a phenotyping algorithm

To highlight the potential of our algorithm for clinical research, we explored the case population identified by our 3+ code algorithm. Cases identified by the 3+ code algorithm were older (median age 69) and had longer record lengths (median 7.3 years) than controls (median age 62, median record length 6.2 years) (Table 3). The percentage of males (53.2%), whites (86.9%), and ever-smokers (54.9%) among COPD cases were higher than among controls (Table 3).

**Table 3.** Demographic and clinical features of COPD cases and controls identified by the 3+ COPD ICD code algorithm, with a minimum floor of 6 months.

| **Characteristic** | **COPD Cases**<br>**N = 28,520** | **Controls**<br>**N = 544,056** | **All SD\***<br>**N = 623,986** |
| --- | --- | --- | --- |
| Median age at last visit (IQR) | 69<br>(60-76) | 62<br>(53-71) | 62<br>(54-72) |
| Sex (%) |  |  |  |
| Female | 13,356<br>(46.8) | 300,402<br>(55.2) | 344,179<br>(55.2) |
| Male | 15,164<br>(53.2) | 243,586<br>(44.8) | 279,738<br>(44.8) |
| Unknown | 0<br>(0.0) | 68<br>(0.0) | 69<br>(0.0) |
| Race (%) |  |  |  |
| White | 24,795<br>(86.9) | 428,349<br>(78.7) | 495,961<br>(79.5) |
| Black | 2,684<br>(9.4) | 45,994<br>(8.5) | 54,547<br>(8.7) |
| Other | 119<br>(0.4) | 6,350<br>(1.2) | 6,908<br>(1.1) |
| Unknown | 922<br>(3.2) | 63,363<br>(11.6) | 66,570<br>(10.7) |
| Smoking status |  |  |  |
| Ever | 15,645<br>(54.9) | 124,295<br>(22.8) | 157,178<br>(25.2) |
| Never | 2,900<br>(10.2) | 213,609<br>(39.3) | 234,382<br>(37.6) |
| Missing | 9,975<br>(35.0) | 206,152<br>(37.9) | 229,249<br>(42.7) |
| Median record length (years)<br>(IQR) | 7.3<br>(3.2-13.0) | 6.2<br>(2.4-11.5) | 6.4<br>(2.5-11.7) |
| Number of respiratory meds**<br>(median, IQR) | 107<br>(32-277) | 16<br>(5-54) | 19<br>(6-67) |
| Top 5 phecodes by frequency<br>(%) |  | 1. |  |
| 1 | Chronic airway obstruction<br>(100) | Hypertension<br>(37.0) | Hypertension<br>(40.9) |
| 2 | Hypertension<br>(77.1) | Essential hypertension<br>(36.2) | Essential hypertension<br>(40.1) |
| 3 | Other symptoms of respiratory<br>system (75.4) | Disorders of lipid<br>metabolism (26.1) | Other symptoms of respiratory<br>system (29.3) |
| 4 | Essential hypertension<br>(75.4) | Hyperlipidemia<br>(26.0) | Disorders of lipid metabolism<br>(28.7) |
| 5 | Tobacco use disorder<br>(63.0) | Other symptoms of<br>respiratory system (24.1) | Hyperlipidemia<br>(28.7) |

\*Individuals with record lengths longer than 180 days.
\*\*Number of respiratory medications includes only mentions of medications on distinct days.

In addition to basic demographic data, we found that cases had a higher median number of respiratory medications per individual than controls (16) (Table 3). We also compared the frequency of phecodes across the two groups.(38–44) In COPD cases, the most frequent phecodes were chronic airway obstruction (100%), hypertension (77.1%), other symptoms of respiratory system (75.4%), essential hypertension (75.4%), and tobacco use disorder (63.0%). In controls, the most frequent phecodes were hypertension (37.0%), essential hypertension (36.2%), disorders of lipoid metabolism (26.1%), hyperlipidemia (26.0%), and other symptoms of respiratory system (24.1%) (Table 3). The most frequent phecodes in the code + regex algorithm cases and PFT cases were similar, with the exception of tobacco use disorder, which was only seen in the five most frequent phecodes for 3+ code cases (Table S3). However, the frequency of the tobacco use disorder phecode was similar in the code + regex cases (68.0%) and in the PFT-defined cases (60.5%) (data not shown). We also describe characteristics of the PFT-defined cases and controls (Table S5).

We performed genetic analyses using single nucleotide polymorphisms from a recent genome-wide association study of COPD by Wain et al.(32) Of the 95 SNPs significantly associated with lung function, 41 were present in our Illumina MEGA array genotyping data. For all but seven of the 41 SNPs present, the confidence intervals from our genetic associations and the confidence intervals from the Wain study overlapped (Figure S3).

## Discussion and Conclusions

Our goal was to develop a phenotyping algorithm for COPD for use in EHR that did not rely on PFTs. While previous studies have also developed phenotyping algorithms for COPD, the population we used in this study is unique. Six of the previously published studies were done in primary care populations in Canada or the United Kingdom.(24, 25, 27, 29–31) Another prior study relied exclusively on insurance claims data.(28) To our knowledge, the only prior study that used an inpatient-based population to electronically phenotype COPD was done by Lacasse et al. However, their primary goal was to determine whether hospital discharge diagnoses of COPD was a valid metric for identifying COPD cases, so they identified cases only and no controls.(26) As a tertiary care center, the VUMC patient population typically represents more complex and severe cases of disease. Furthermore, the Southeast United States, where VUMC is located, has a higher smoking prevalence and COPD prevalence than the national average.(1, 47) Implementation of a COPD phenotyping algorithm in clinical centers where COPD prevalence is among the nation’s highest represents an opportunity to utilize our electronic health systems to address a leading cause of morbidity and mortality.

We also sought to develop an algorithm that was easy to implement so it could be deployed across different health care systems to enable clinical research and pragmatic clinical trials. In addition to calculating clinical validity metrics, we considered demographic and clinical characteristics of the resulting COPD case and control groups when selecting our algorithm. While performance metrics are valuable, EHR present unique challenges that necessitate a more nuanced approach. Data missingness is a common problem in EHR-based research.(48– 56) This is particularly true in a tertiary referral center such as VUMC, where patients often receive routine clinical care at other institutions. We found that adding more stringent criteria for cases did improve our PPV, but did not greatly impact the clinical characteristics of the case sets identified (Table S4). The strength of association between COPD and established risk factors such as age, sex, and smoking did not differ between our high PPV code + regex algorithm cases and the less stringent 3+ code algorithm cases. The genetic profile of the cases identified by our 3+ code algorithm is also consistent with previous research (Figure S3).(32) By requiring more clinical data, our code + regex algorithm biased our case sample to individuals with more severe disease, since sicker patients typically have denser documentation and more complete data in EHR.(56)

We recognize that in some research settings, a more stringent case definition and higher PPV may be more appropriate. In such settings, our code + regex algorithm may be a better choice, since it can achieve a PPV of up to 86.5% (Table 2). For facilities that lack the phenotyping resources to extract more complex clinical data such as oxygen use, we also tested a case definition requiring 5+ ICD codes only. This algorithm had a higher PPV (70.6%) in our chart review validation set than our 3+ code-only algorithm (57.1%). The specificity of the 5+ code-only algorithm was also higher (86.4%) compared to the 3+ code-only algorithm (76.0%). The NPV remained the same, and only the sensitivity decreased slightly (97.3% in the 5+ code-only algorithm vs. 97.6% in the 3+ code-only algorithm) (data not shown). The same pattern in performance changes was seen in the PFT validation set (data not shown). This relatively simple change to our 3+ code-only algorithm can be used in settings where more advanced phenotyping is not possible.

For all our algorithms, we found that the performance was higher in the chart review validation set than in the PFT set. This is likely due to the enrichment of respiratory disease in the PFT set. PFTs are not performed for routine screening, so individuals referred for PFT typically have some respiratory compromise.(57) Many of the variables we included in our algorithms are present in individuals with other respiratory conditions. A few of the individuals classified as cases by our algorithm had PFTs that were not consistent with the GOLD definition of COPD.(8) This may be due to an absence of temporal data in our study. We did not require individuals to have received PFTs and ICD codes for COPD in any particular order or time frame. Therefore, individuals labeled as controls in the PFT set may have received PFTs for early respiratory decline that did not meet the official definition for COPD at the time. It is possible that these individuals experienced continued lung function decline that eventually progressed to COPD, but never received follow-up PFTs at VUMC. The overlap between COPD and other respiratory conditions and the high prevalence of respiratory disease in our PFT set likely explain the reduced algorithm performance in the PFT set.

This study has several limitations. As previously mentioned, the use of EHR in biomedical research has inherent challenges due to inconsistent documentation, missing data, and inaccuracies.(48–56) Tertiary care EHR data often have sicker patients with denser clinical documentation and more complete data than other hospital settings.(56) COPD cases identified by our algorithm have more severe COPD based on FEV_1_ and FEV_1_/FVC measurements (Figure S2). Our algorithm relied heavily on ICD code information, which can be inaccurate, particularly for secondary research use.(58, 59) However, our analyses of known COPD risk factors and previous genetic associations suggest that the population identified by our algorithm has similar epidemiologic and genetic characteristics to previously studied COPD populations.(60) Furthermore, we present a second algorithm, the code + regex algorithm, that includes oxygen use on the problem list in addition to ICD codes. This approach has been previously demonstrated to improve phenotyping accuracy.(58) Another limitation is the predominance of individuals of European descent in our study population. This may limit generalizability to other populations. Replication in other EHR systems with greater diversity is needed to address this.

A key advantage to our case population is the wealth of clinical information contained within the EHR, which can be leveraged for clinical research. The addition of a linked DNA repository provides unique opportunities for COPD genetics research. Application of our phenotyping algorithm allowed us to identify a large population with COPD for research, without the additional time and cost investments typically required to build epidemiologic cohorts.

Overall, our study demonstrates that phenotyping algorithms for COPD can be successfully implemented in EHR in a tertiary hospital setting. We present several algorithms with different clinical validity metrics, as we recognize the best algorithm may vary depending on the research question. Use of COPD phenotyping algorithms can quickly and easily identify large cohorts for clinical research studies within EHR, which will facilitate accelerated scientific discoveries and precision medicine opportunities for this devastating disease.

## Acknowledgements

This work was supported in part by NIGMS (T32GM007347 and T32GM080178), NHLBI (F30HL140756) and the Vanderbilt CTSA grant UL1TR002243 from NCATS/NIH.

