## Supplemental Tables and Figures for "A Clinical Phenotyping Algorithm to Identify Cases of Chronic Obstructive Pulmonary Disease in Electronic Health Records"

**Table S1**. ICD codes and semantic phrases considered in algorithm development

| **Inclusion Criteria for Cases** | **Additional Exclusion Criteria for Controls** |
|---|---|
| ICD-9 Codes | ICD-9 Codes |
| 491.xx: Chronic bronchitis | 493.xx: Asthma |
| 492.xx: Emphysema | 516.31: Idiopathic pulmonary fibrosis |
| 496.xx: Chronic airway obstruction, not elsewhere classified | 135.xx: Sarcoidosis |
| 162.xx: Neoplasm of the trachea, bronchus, and lung | |
| ICD-10 Codes | ICD-10 Codes |
| J41.xx: Simple and mucopurulent chronic bronchitis | J45.xx: Asthma |
| J42.xx: Unspecified chronic bronchitis | J84.112: Idiopathic pulmonary fibrosis |
| J43.xx: Emphysema | D86.xx: Sarcoidosis |
| J44.xx: Other chronic obstructive pulmonary disease | |
| C34.xx: Malignant neoplasm of bronchus and lung | |
| Problem list text | |
| COPD | |
| Chronic obstructive pulmonary disease | |
| Chronic obstructive lung disease | |
| Emphysema | |
| Chronic bronchitis | |
| Reactive airway(s) disease | |
| Exacerbation | |
| Oxygen | |
| O2 | |
| Radiology report text | |
| Emphysema (excluding subcutaneous emphysema) | |
| Emphysematous changes | |
| Respiratory medications | |
| Accuneb | |
| Aclidinium | |

| |
|---|
| Advair |
| Albuterol |
| Alvesco |
| Anoro |
| Arcapta |
| Arformoterol |
| Asmanex |
| Atrovent |
| Beclomethasone |
| Beclovent |
| Breo |
| Brovana |
| Budesonide |
| Ciclesonide |
| Combivent |
| Daliresp |
| Dulera |
| Duoneb |
| Entocort |
| Flovent |
| Fluticasone |
| Foradil |
| Glycopyrrolate |
| Indaceterol |
| Ipratropium |
| Lasma |
| Levalbuterol |
| Mometasone |
| Olodaterol |
| Perforomist |
| ProAir |
| Proventil |
| Pulmicort |
| QVar |
| Robinul |
| Roflumilast |
| Salbutamol |

| |
|---|
| Salmeterol |
| Serevent |
| Singulair |
| Spiriva |
| Symbicort |
| Theo |
| Theochron |
| Theodur |
| Theophylline |
| Tiotropium |
| Tudorza |
| Umeclidinium |
| Uniphyl |
| Vanceril |
| Ventolin |
| Vilanterol |
| Xopenox |

**Table S2.** Numbers of cases, controls, indeterminates, and excludes identified among adults over 45 years of age in the Synthetic Derivative (N = 1,008,552).

| Case definition* | Cases, N | Control, N | Indeterminate, N | Exclude, N |
|---|---|---|---|---|
| 3+ ICD codes | 28,520 | 544,056 | 23,091 | 28,319 |
| ICD code+reg-expr | 12,622 | 544,056 | 38,989 | 28,319 |
| PFT | 1,873 | 11,203 | NA | NA |

PFT = pulmonary function test, reg ex = regular expression

*All case definitions also include the requirement that cases have records longer than 180 days.

**Table S3**. Demographic characteristics of COPD cases identified by PFT, 3+ ICD codes, and Code + regex algorithm.

| **Variable** | **PFT cases*<br>N = 1,873** | **3+ COPD ICD Cases<br>N = 28,520** | **Code+Regex Cases<br>N = 12,622** |
|---|---|---|---|
| Median age (IQR) | 70<br>(62-76) | 69<br>(60-76) | 70<br>(62-77) |
| Sex (%) | | | |
| Female | 911<br>(48.6) | 13,356<br>(46.8) | 6,163<br>(48.8) |
| Male | 962<br>(51.4) | 15,164<br>(53.2) | 6,459<br>(51.2) |
| Unknown | 0<br>(0.0) | 0<br>(0.0) | 0<br>(0.0) |
| Race (%) | | | |
| White | 1,701<br>(90.8) | 24,795<br>(86.9) | 11,012<br>(87.2) |
| Black | 135<br>(7.2) | 2,684<br>(9.4) | 1,249<br>(9.9) |
| Other | 15<br>(0.8) | 119<br>(0.4) | 46<br>(0.4) |
| Unknown | 22<br>(1.2) | 922<br>(3.2) | 315<br>(2.5) |
| Smoking status | | | |
| Ever | 1,464<br>(78.2) | 15,645<br>(54.9) | 7,548<br>(59.8) |
| Never | 402<br>(21.5) | 2,900<br>(10.2) | 1,034<br>(8.2) |
| Missing | 7<br>(0.4) | 9,975<br>(35.0) | 4,040<br>(32.0) |
| Median record length (years) (IQR) | 10.3<br>(5.1-16.1) | 7.3<br>(3.2-13.0) | 8.3<br>(4.0-13.8) |
| Number of respiratory meds** (median, IQR) | 188<br>(72-385) | 107<br>(32-277) | 170<br>(56-387) |
| Top 5 phecodes by frequency (%) | | | |

| | | | |
|---|---|---|---|
| 1 | Other symptoms of respiratory system (88.7) | Chronic airway obstruction (100) | Chronic airway obstruction (100) |
| 2 | Chronic airway obstruction (78.0) | Hypertension (77.1) | Other symptoms of respiratory system (86.1) |
| 3 | Shortness of breath (71.8) | Other symptoms of respiratory system (75.4) | Hypertension (80.3) |
| 4 | Hypertension (71.8) | Essential hypertension (75.4) | Essential hypertension (78.9) |
| 5 | Essential hypertension (70.7) | Tobacco use disorder (63.0) | Shortness of breath (73.5) |

*Pulmonary function testing (PFT) cases are defined as individuals with a post-bronchodilator $FEV_1$/FVC ratio < 0.7.

**Number of respiratory medications includes only mentions of medications on distinct days.

**Table S4.** Association of known clinical risk factors with COPD in each algorithm set and in the PFT set.

| | 3+ code only | | Code + regex | | PFTs | |
|---|---|---|---|---|---|---|
| **Risk factor** | **OR** | **95% CI** | **OR** | **95% CI** | **OR** | **95% CI** |
| Age | 1.04 | 1.04-1.04 | 1.05 | 1.05-1.05 | 1.04 | 1.03-1.04 |
| Sex | 1.40 | 1.37-1.43 | 1.27 | 1.23-1.31 | 1.22 | 1.11-1.34 |
| Ever smoking | 9.22 | 8.87-9.57 | 12.36 | 11.61-13.16 | 3.36 | 3.00-3.75 |

OR: odds ratio; 95% CI: 95% confidence interval

**Table S5**. Comparison of demographics and clinical features for cases (post-bronchodilator $FEV_1$/FVC < 0.7) and controls identified by PFTs.

| **Characteristic** | **Cases*<br>N = 1,873** | **Controls*<br>N = 11,203** | **All PFT*<br>N = 13,076** |
|---|---|---|---|
| Median age (IQR) | 70<br>(62-76) | 65<br>(57-72) | 65<br>(57-73) |
| Sex (%) | | | |
| Female | 911<br>(48.6) | 6,023<br>(53.8) | 6,934<br>(53.0) |
| Male | 962<br>(51.4) | 5,179<br>(46.2) | 6,141<br>(47.0) |
| Unknown | 0<br>(0.0) | 1<br>(0.0) | 1<br>(0.0) |
| Race (%) | | | |
| White | 1,701<br>(90.8) | 9,717<br>(86.7) | 11,418<br>(87.3) |
| Black | 135<br>(7.2) | 1,201<br>(10.7) | 1,336<br>(10.2) |
| Other | 15<br>(0.8) | 162<br>(1.4) | 177<br>(1.4) |
| Unknown | 22<br>(1.2) | 123<br>(1.1) | 145<br>(1.1) |
| Smoking status | | | |
| Ever | 1,464<br>(78.2) | 5,779<br>(51.6) | 7,243<br>(55.4) |
| Never | 402<br>(21.5) | 5,359<br>(47.8) | 5,761<br>(44.1) |
| Missing | 7<br>(0.4) | 65<br>(0.6) | 72<br>(0.5) |
| Median record length (years) (IQR) | 10.3<br>(5.1-16.1) | 9.5<br>(4.5-15.5) | 9.6<br>(4.6-15.6) |
| Number of respiratory meds** (median, IQR) | 188<br>(72-385) | 168<br>(66-372) | 171<br>(67-374) |
| Top 5 phecodes by frequency (%) | | | |
| 1 | Other symptoms of respiratory system (88.7) | Other symptoms of respiratory system (84.5) | Other symptoms of respiratory system (85.1) |

| | | | |
|---|---|---|---|
| 2 | Chronic airway obstruction (78.0) | Hypertension (66.7) | Hypertension (67.1) |
| 3 | Shortness of breath (71.8) | Essential hypertension (65.6) | Essential hypertension (66.3) |
| 4 | Hypertension (71.8) | Shortness of breath (64.1) | Shortness of breath (65.2) |
| 5 | Essential hypertension (70.7) | Diseases of esophagus (54.9) | Disorders of lipoid metabolism (55.2) |

*Cases, controls, and all PFT data sets were restricted to individuals with record lengths greater than 180 days to be comparable to those identified by the phenotyping algorithm.
**Number of respiratory medications includes only mentions of medications on distinct days.

**Figure S1.** Phenotyping algorithm with highest PPV in the chart review validation set.

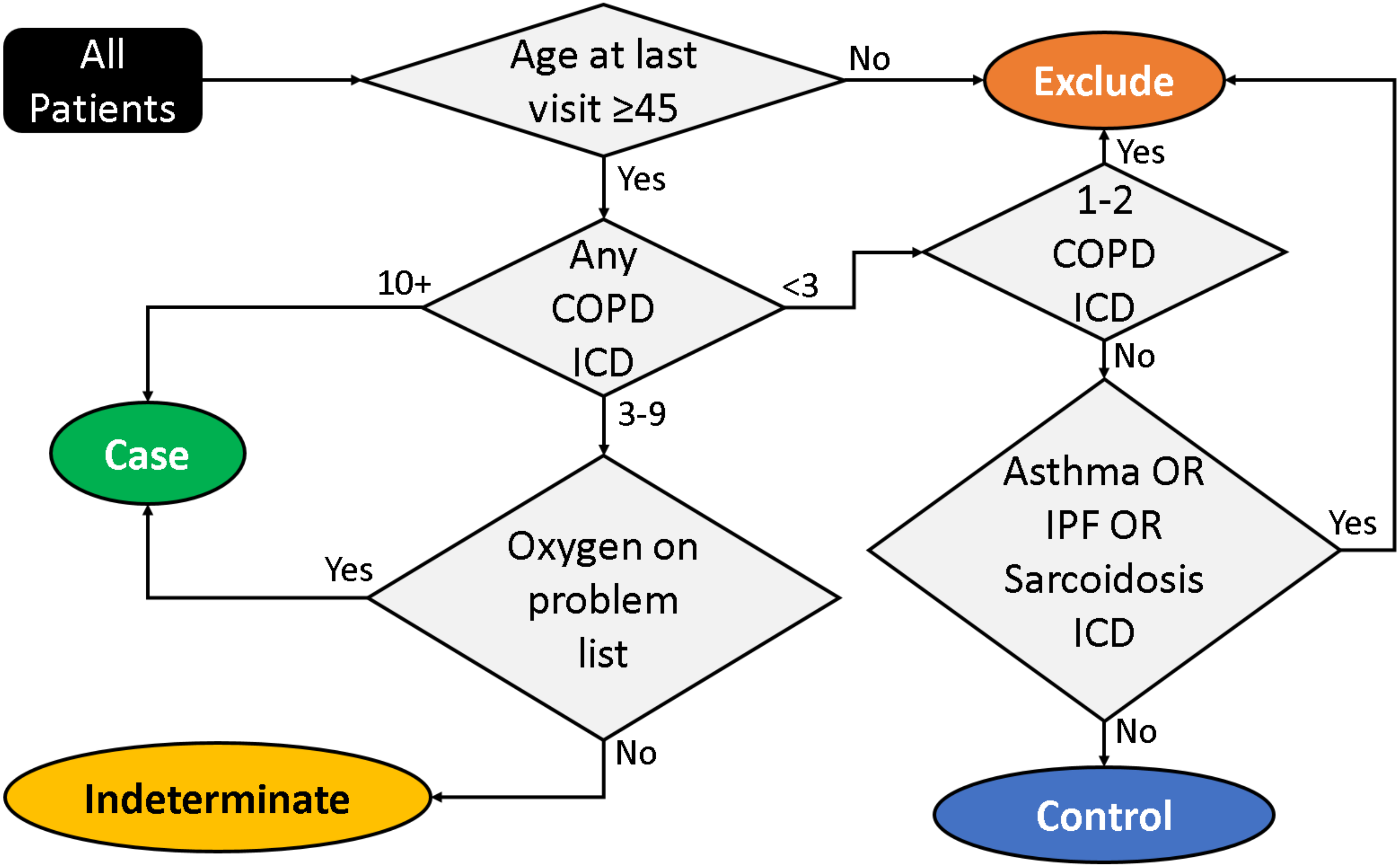

**Figure S2.** Comparison of PFT-defined cases with 3+ ICD codes for COPD and those with fewer than three ICD codes for COPD among individuals with record lengths greater than 180 days. A) Forced expiratory volume in 1 second, pre-bronchodilator. B) Forced expiratory volume in 1 second, post-bronchodilator. C) Forced expiratory volume in 1 second/forced vital capacity, pre-bronchodilator. D) Forced expiratory volume in 1 second/forced vital capacity, post-bronchodilator.

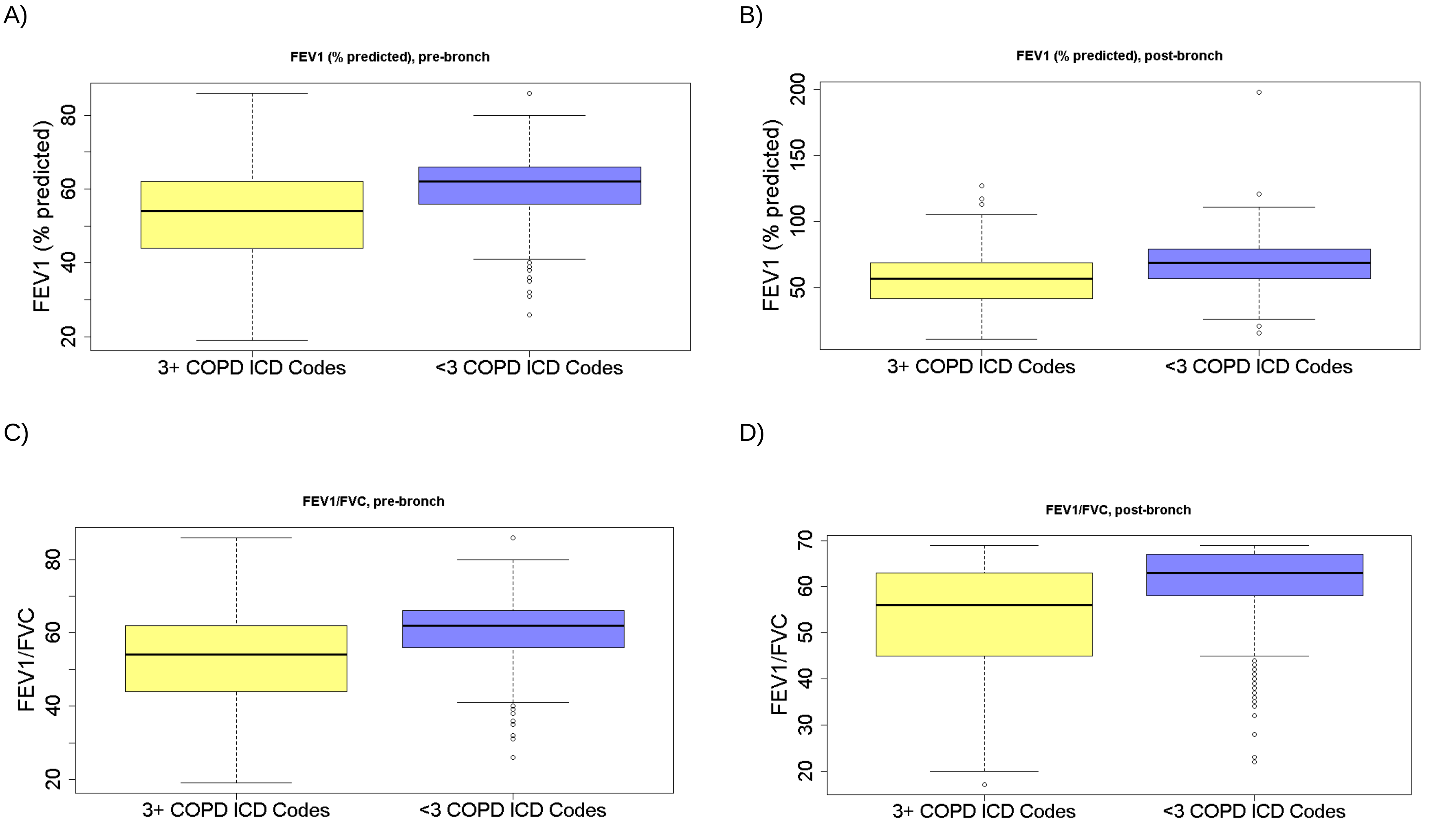

**Figure S3**. Odds ratios from genetic associations of COPD cases (3+ code) and controls in BioVU compared with associations of select variants identified in a prior GWAS of COPD by Wain et al. (32)

| CHR | BP | SNP |
|---|---|---|
| 1 | 17306675 | rs2284746 |
| 1 | 40035686 | rs17513135 |
| 1 | 92068967 | rs1192404 |
| 1 | 150586971 | rs6681426 |
| 1 | 218860068 | rs993925 |
| 1 | 219963088 | rs4328080 |
| 2 | 18309132 | rs62126408 |
| 2 | 218683154 | rs2571445 |
| 2 | 229502503 | rs10498230 |
| 2 | 239316560 | rs61332075 |
| 2 | 239877148 | rs12477314 |
| 3 | 67452043 | rs1490265 |
| 3 | 127991527 | rs2811415 |
| 4 | 89870964 | rs2045517 |
| 4 | 106688904 | rs10516526 |
| 4 | 106819053 | rs34712979 |
| 5 | 52195033 | rs1551943 |
| 5 | 131788334 | rs7713065 |
| 5 | 147856392 | rs7715901 |
| 5 | 156810072 | rs10515750 |
| 6 | 27459923 | rs34864796 |
| 6 | 109270656 | rs2768551 |
| 6 | 142838173 | rs148274477 |
| 9 | 4124377 | rs7872188 |
| 9 | 98204792 | rs16909859 |
| 9 | 139102831 | rs10858246 |
| 9 | 139257411 | rs10870202 |
| 10 | 69957350 | rs7095607 |
| 10 | 78312002 | rs2637254 |
| 11 | 45250732 | rs2863171 |
| 12 | 28689514 | rs2348418 |
| 12 | 57527283 | rs11172113 |
| 12 | 65824670 | rs1494502 |
| 12 | 95554771 | rs113745635 |
| 12 | 115201436 | rs10850377 |
| 12 | 115500691 | rs35506 |
| 15 | 41977690 | rs72724130 |
| 16 | 78187138 | rs1079572 |
| 17 | 36886828 | rs11658500 |
| 21 | 35690499 | rs2834440 |
| 22 | 18450287 | rs11704827 |

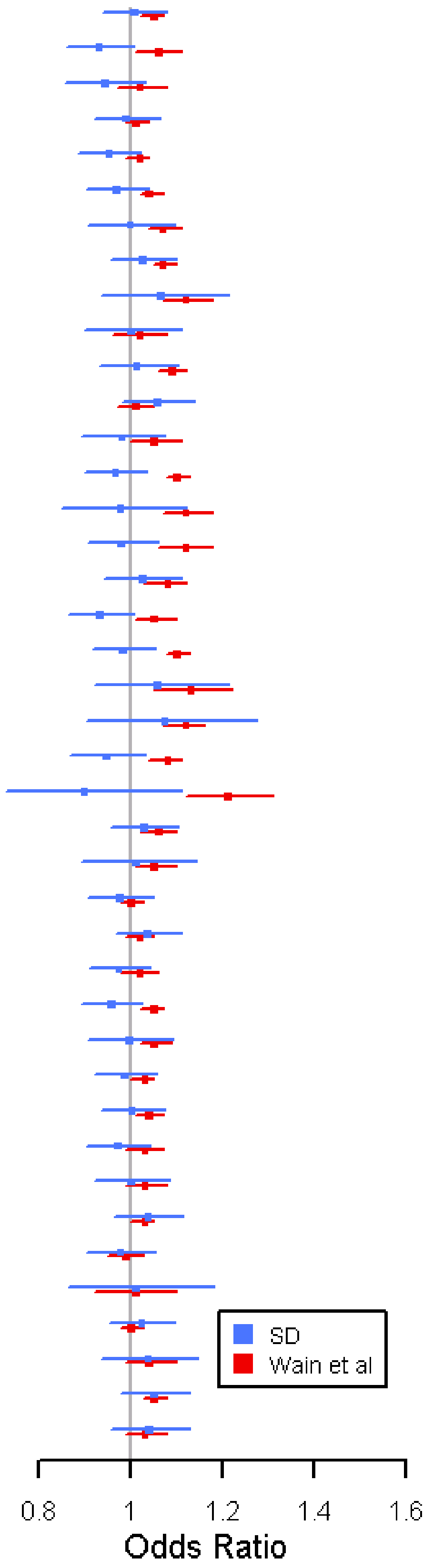
